# LRMD: Reference-Free Misassembly Detection Based on Multiple Features from Long-Read Alignments

**DOI:** 10.1101/2025.11.07.686952

**Authors:** Xianfeng Shi, Fan Nie, Jianxin Wang

## Abstract

Genome assembly serves as the cornerstone of genomics research, with the detection of misassembly playing a crucial role in downstream analyses. Reference-free methods for misassembly detection, leveraging read alignments, enable us to circumvent the need for high-quality reference genomes and broaden their applicability. However, existing methods struggle to effectively utilize alignment data, leading to a noticeable deficiency in sensitivity for detecting misassemblies. We introduce LRMD, a novel reference-free tool for misassembly detection. LRMD integrates depth, clipping, and read pileup information derived from long-read-to-assembly alignments to significantly enhance sensitivity in identifying misassemblies. Experimental evaluations on both simulated and real datasets demonstrate that LRMD consistently outperforms existing tools in terms of sensitivity and F1-score. Notably, its results are closest to the reference-based evaluation tool QUAST. As an evaluation tool, LRMD also outputs metrics such as base quality, assembly size, contig N50, and others. LRMD is public available at http://github.com/sxfss/LRMD.

## 1 INTRODUCTION

High-quality genome assemblies form the cornerstone of research into species evolution, gene expression, population genetics, and more. The advent of third-generation long-read sequencing technologies [1, 2] has spurred the development of numerous assembly tools [3–8], resulting in significant advancements in assembly quality [9–11], particularly in terms of assembly continuity. However, due to the complexity of genomic regions such as repeats and centromeres, coupled with relatively high sequencing errors inherent in long reads, assembly outcomes still contain a notable number of errors. These errors can severely impact downstream analyses [12–14], underscoring the critical need for thorough and effective evaluation of genome assemblies [9, 15, 16].

Several metrics are employed to assess assembly quality from different perspectives. Contig N50 is widely used to gauge assembly continuity but overlooks assembly errors within contigs. . GCI [17] privdes an another score to quantify the continuity of high-quality genomes, which takes potential assembly errors into account. Base-level accuracy assesses small-scale errors like mismatches and small indels. Gene completeness measured by BUSCO [18] counts conserved genes to evaluate assembly comprehensiveness. The number of misassemblies identifies structural errors due to incorrect read connections, encompassing deletions, insertions, inversions, duplications, translocations, and other types. Unlike base-level errors, misassemblies are often challenging to correct with polishing tools and profoundly impact downstream genomics studies, including evolutionary analysis [19]. Structural variant (SV) detection [20–22] is a process similar to misassembly detection. However, while SV detection focuses on discovering genomic differences, misassembly detection is concerned with identifying errors resulting from incorrect assembly.

Currently, accurately detecting misassemblies remains a significant challenge in genome assembly assessment. Two primary types of methods are used for structural misassembly detection: reference-based and reference-free approaches. Reference-based methods typically involve aligning the assembled genome to a known reference sequence to identify inconsistencies. Tools like QUAST [23, 24] and GAGE [25] evaluate assembly quality by comparing against reference sequences, categorizing discrepancies such as inversions, relocations, and translocations. These methods theoretically offer comprehensive detection capabilities when a high-quality reference genome is available. However, many species lack such a reference, limiting the applicability of these methods. Additionally, reference-based approaches may struggle to differentiate between genuine misassemblies and natural genetic variations arising from individual diversity differences between reference and target genomes, further constraining their practical utility.

Reference-free methods for misassembly detection, which assess alignments from reads to assemblies, are widely applicable. Tools designed for short reads [26–29] detect misassemblies by analyzing factors such as inconsistent coverage in alignments and the distribution of paired-end reads. While long reads offer advantages in detecting larger misassemblies due to their length, their higher sequencing error rates and uneven coverage can impact detection performance. Inspector [30] identifies misassembly signals from alignments and employs clustering with variable window sizes to filter out false-positive candidates. CRAQ [31] identifies regional and structural assembly errors by utilizing clipped alignment information. GAEP [32] detects potential misassembly breakpoints by scanning clips within alignments. These existing methods primarily rely on extracting some obvious information such as clipping information to detect misassemblies. While these methods achieve high accuracy, their sensitivity can be limited.

In this study, we introduce LRMD, a novel reference-free tool for detecting misassemblies. LRMD leverages multiple features extracted from long-read-to-assembly alignments to identify regions potentially affected by misassemblies. An advanced window filtering algorithm enhances the accuracy of detection. Additionally, LRMD provides output metrics including base quality and assembly continuity (N50). Through benchmarking on simulated and real datasets, LRMD demonstrates significantly improved sensitivity in detecting misassemblies. These findings underscore LRMD’s effectiveness in leveraging alignment data for robust misassembly detection.

## 2 METHODS

### 2.1 Overview of LRMD

LRMD is a practical tool designed to detect misassemblies by analyzing alignments between long reads and assemblies. It operates through three main steps, outlined in **Figure 1**: (1) mapping long reads to the assembly; (2) identifying regions suspected of containing misassemblies; and (3) applying a filtering process to reduce false positives and enhance accuracy. In addition to misassembly detection, LRMD offers several output metrics such as base quality, number of contigs, total assembly size, largest contig size, and N50 contig length. These features make LRMD a comprehensive tool for assessing and refining genome assemblies based on long-read data.

**Figure 1:**
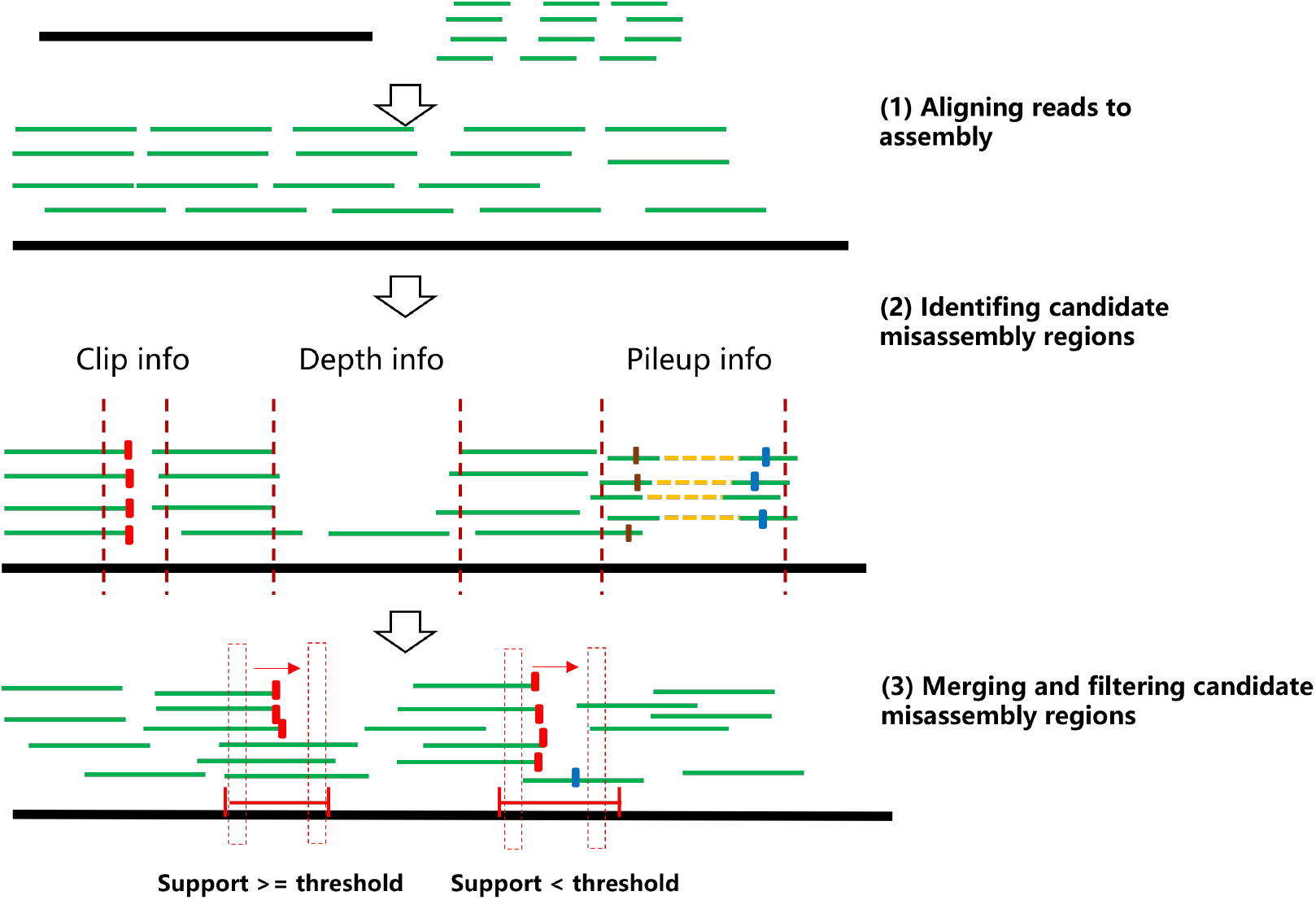
LRMD misassembly detection workflow.

### 2.2 Read alignments

In this study, Minimap2 [33] is utilized for aligning long reads to assemblies. Specifically, the parameters “-a -x map-pb” are applied for PacBio CLR reads, “-a -x map-hifi” for PacBio HiFi reads, and “-a -x map-ont” for Nanopore reads. Once SAM files are generated by minimap2, samtools is employed to convert them into BAM format, followed by sorting and indexing for subsequent analysis.

### 2.3 Identifying candidate misassembly regions

Based on practical observations, regions exhibiting misassembly often display notable characteristics such as drastic changes in coverage depth, frequent read clips, and an abundance of mismatches and indels (referred to as noisy alignment regions), as illustrated in **Figure 2**. To identify candidate misassembly regions, LRMD employs a sliding window approach, typically set at a default size of 400 bp, to detect and highlight these features.

**Figure 2:**
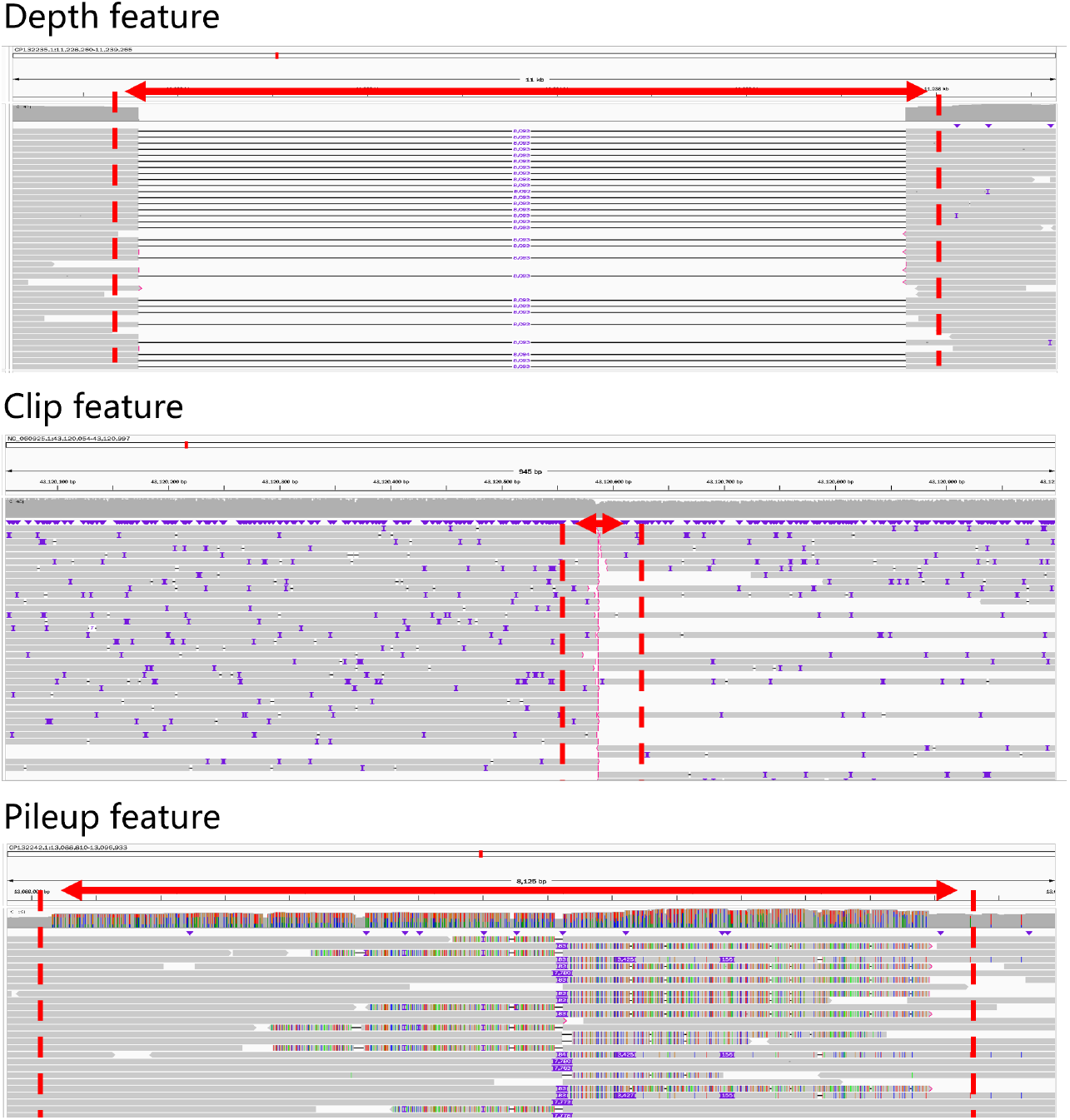
Examples of features used by LRMD.

#### (1) Depth information

The average coverage depth *d*_*w*_ of each sliding window is calculated. The window is considered as a candidate misassembly region, if *d*_*w*_ satisfies the condition: *d*_*w*_ *< d*_*g*_ × *p*_*l*_ or *d*_*w*_ *> d*_*g*_ × *p*_*u*_, Here, *d*_*g*_ is the global average coverage depth, while *p*_*l*_ and *p*_*u*_ is user-defined parameters (default values are 0.15 and 0.75, respectively).

#### (2) Read clips information

A clip refers to a substantial discrepancy between the read and the reference sequence within a region. After filtering out unmapped reads, secondary alignments and low-quality alignments, LRMD scans the BAM file and tallies the number of clips *c*_*w*_ (where the clip *length* ≥ 500) within each sliding window. If *c*_*w*_ meets the condition: *c*_*w*_ ≥ *p*_*w*_ × *d*_*g*_, the window is considered as a candidate misassembly regions. Here, *p*_*w*_ is a user-defined parameter with a default value is 0.35, and *d*_*g*_ denotes the global average coverage depth.

#### (3) Pileup information

A pileup file generated by the samtools mpileup command provides counts of matches, mismatches, and indels at each position of the assembly. LRMD calculates the percentages of matches (*p*_*match*_ ), mismatches (*p*_*mismatch*_ ) and indels (*p*_*indel*_ ) within each sliding window based on this information. *p*_*indel*_ is calculated as the sum of the lengths of INDELs, rather than the number of INDELs, divided by the length of the window. The related sliding window is considered as a candidate misassembly region. if any of the following conditions are met:

- *p*_*match*_ ≤ *th*_*match*_
- *p*_*mismatch*_ ≥ *th*_*mismatch*_
- *p*_*indel*_ ≥ *th*_*indel*_

Here, *th*_*match*_, *th*_*mismatch*_ and *th*_*indel*_ are user-defined thresholds (default values are 0.9, 0.1, 0.2 respectively for Nanopore data; 0.95, 0.05, 0.15 respectively for PacBio HiFi data).

### 2.4 Merging and filtering candidate misassembly regions

After identifying candidate misassembly regions using the steps described above, LRMD proceeds to merge overlapping regions or regions that are within 2000 bp of each other into larger contiguous regions. Following merging, LRMD applies a sliding window approach with a length of 6000 bp for PacBio HiFi reads, and 10000 bp for Nanopore and PacBio CLR reads, sliding over each candidate mis-assembly region. LRMD filters out candidate misassembly regions if each sliding window within these regions contains sufficient high-quality alignments to cover the entire window. The high-quality alignment is the primary alignment, where the cumulative length of large insertions (>=30bp) or large deletions (>=30bp) does not exceed 100. The number of high-quality alignments (*N*_*hq*_ ) needs to meet the following conditions: *N*_*hq*_ *>*= max (*N*_*th*_, 0.4 ∗ *N*_*w*_ ), where *N*_*w*_ is the number of reads across the sliding windows and *N*_*th*_ is a user-defined threshold (default values are 3 for Nanopore data and for PacBio HiFi data).

### 2.5 Base quality evaluation

The base quality value (QV) assesses the incidence of small scale mis-assembly in the assembly from an overall perspective. The pileup file generated by samtools provides counts of matches (*n*_*match*_ ), mismatches (*n*_*mismatch*_ ), and indels (*n*_*indel*_ ) at each position of the assembly. If the condition: 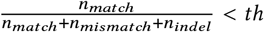 is met, we consider that the base at the corresponding position in the assembly is incorrect. *th* is a user-defined threshold (default value is 0.7). We count the number of all bases (*N*_*total*_ ) and the bases considered as error (*N*_*error*_ ) in the assembly. QV can be calculated as 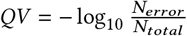.

### 2.6 Simulated datasets and evaluation

#### 2.6.1 Simulated datasets

To evaluate the performance of LRMD, we generated three simulated assemblies using reference genomes CHM13, *Oryza sativa* (*O. sativa*) and GRCH38 as templates, as show in **Table 1**. The simulation method was adapted from GAEP [32], which can introduce five types of misassemblies (insertion, delection, inversion, expansion and collapse). Those misassemblies vary in length from 1000 bp to 50,000 bp. Specifically, the simulated assemblies resulted in 4298, 826 and 2238 misassemblies for GRCH38, CHM13, and *O. sativa* templates, respectively.

**Table 1:** Detailed information of the real datasets.

| Species | Data Type | Download Link |
| --- | --- | --- |
| CHM13 | fastq | CHM13 rel6, SRX789768 |
|  | reference | T2T-CHM13v2.0 |
| *Oryza sativa* | fastq | SRX20987366, SRX20987367 |
|  | reference | GCA_034140825.1 |
| GRCH38 | reference | GRCh38.p14 |

We used the read simulation tool pbsim3 [34] and the data simulator pbccs(https://github.com/PacificBiosciences/ccs?tab=readme-ov-file) provided by Pacific Biotechnology to generate our simulated nanopore reads and PacBio reads. For simulation, generate PacBio HiFi and Nanopore reads from the simulated assemblies of CHM13 and *O. sativa*, and PacBio CLR reads from the simulated assembly of GRCH38. Detailed simulation commands can be found in **Supplementary A**.

#### 2.6.2 Evaluation

Due to the deviation in the positions between the misassemblies identified by the detection methods and those simulated, we consider them to be the same misassembly if they overlap or are within a distance of 2000 bp. Using these criteria, we calculate the precision, recall, and F1 score for each detection method on simulated datasets.

### 2.7 Read datasets and evaluation

#### 2.7.1 Read datasets

Multiple real assemblies were utilized to assess LRMD’s performance. We employed two PacBio HiFi datasets from CHM13 and O. sativa, as well as two Nanopore datasets from CHM13 and O. sativa for assembly, as show in **Table 1**. To standardize the sequencing data, reads were down-sampled to 40X coverage using rasusa, whenever the coverage exceeded this threshold. For Nanopore reads, assemblies were conducted separately using flye [6] and wtdbg2 [5]. PacBio HiFi reads were assembled using flye and hifiasm [7], also separately.

#### 2.7.2 Evaluation

The Jaccard coefficient is used to assess the similarity of misassemblies identified by different tools. To mitigate impact from shorter low-quality contigs, detection results from contigs shorter than 40kb were excluded from the analysis. Additionally, we observed that QUAST tended to split several large misassemblies into numerous smaller ones in certain regions, introducing bias in the evaluation. To address this, we consolidated QUAST’s detection results based on proximity. Specifically, we merged misassemblies that were within a 5kb distance of each other. We consider two misassemblies to be the same if they overlap or are within a distance of 1000 bp. Due to one misassembly detected by one tool possibly corresponding to multiple misassemblies detected by other tools, we calculate the Jaccard coefficient based on the proportion of misassemblies rather than their count. The Jaccard coefficient is calculated by 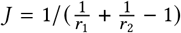, where *r*_1_ and *r*_2_ are the proportions of misassemblies detected by one tool that are contained within those detected by another tool.

## 3 RESULTS

### 3.1 Benchmark on simulated datasets

To benchmark the performance of LRMD, we compared it with three other reference-free assembly evaluators: GAEP, CRAG, and Inspector, as well as the reference-based assembly evaluator QUAST, using simulated datasets derived from three reference genomes: *O. sativa*, CHM13, and GRCH38 (details in **Methods**). We simulated PacBio HiFi reads and Nanopore reads from *O. sativa* and CHM13, and PacBio CLR reads from GRCH38. The simulated assemblies comprised 826 misassemblies for *O. sativa*, 4298 for CHM13, and 2238 for GRCH38 (see **Table 2**). LRMD demonstrated the closest detection of the actual number of assembly errors. To accommodate potential differences where methods might split one misassembly into two or combine two misassemblies into one, we allowed for a certain offset (2000 bp) when calculating the precision and recall of these methods (see **Methods**).

**Table 2:** The number of misassemblies detected by the different tools on the simulated datasets.

| Dataset | Simulate<sup>1</sup> | GAEP | CRAQ | Inspector | QUAST | LRMD |
| --- | --- | --- | --- | --- | --- | --- |
| O. sativa, ONT | 826 | 1353 | 1104 | 2332 | 2083 | 882 |
| CHM13, ONT | 4298 | 5957 | 4718 | 10895 | 9814 | 4653 |
| O. sativa, HiFi | 826 | 1351 | 2109 | 2375 | 2083 | 896 |
| CHM13, HiFi | 4298 | 6094 | 10253 | 11215 | 9814 | 4662 |
| GRCH38, CLR | 2238 | 3587 | 1756 | 5605 | 4725 | 2588 |

<sup>1</sup> Simulate represents the number of simulated misassemblies.
<sup>2</sup> QUAST is a reference-based tools, and the others are reference-free tools.

In **Figure 3**, LRMD outperforms other reference-free methods such as CRAQ, GAEP, and Inspector across Nanopore, PacBio HiFi, and PacBio CLR datasets in terms of F1 score. While Inspector achieves 100% precision on all datasets, its recall rate is lower, ranging from 41.2% to 48.2%. CRAQ shows similar high precision but also suffers from low recall. LRMD and GAEP strike a better balance between precision and recall. LRMD particularly excels in recall on simulated datasets, achieving 96.7% to 99.1%, surpassing the other methods. This is attributed to LRMD’s innovative approach of integrating multiple features for misassembly detection. LRMD also demonstrates robust detection precision across Nanopore, PacBio HiFi, and PacBio CLR datasets, indicating consistent performance irrespective of data type. In theory, reference-based tools like QUAST should perform optimally with a perfect reference sequence. However, on the O. sativa dataset, QUAST achieves slightly less than 90% precision. Analysis reveals issues with sequence alignment in the 0-4 MB interval of the CP132243.1 chromosome, leading to numerous false positives in QUAST evaluation and hence lower precision.

**Figure 3:**
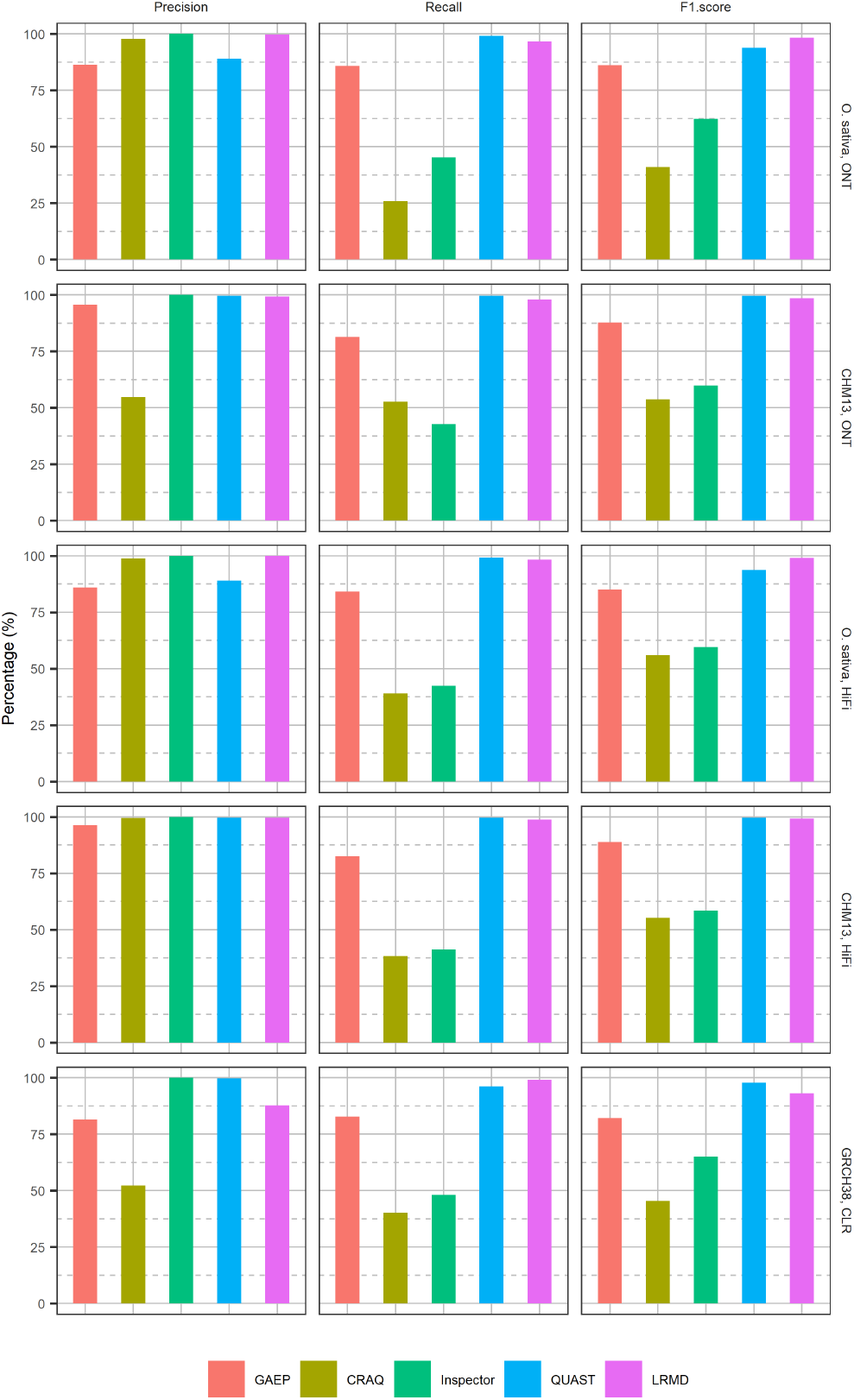
Precison, recall and F1 score of the tools on the simulated datasets.

Overall, LRMD’s comprehensive approach to misassembly detection, leveraging advanced feature integration and achieving high recall rates, underscores its effectiveness in genome assembly evaluation across different sequencing technologies.

### 3.2 Ablation Experiment on Simulated Chm13 PacBio HiFi dataset

LRMD integrates various features from long-read-to-assembly alignments to enhance detection sensibility. To assess the impact of each feature, we conducted an ablation experiment using simulated CHM13 PacBio HiFi dataset. The experiment design involved evaluating each feature type individually, then combinations of two types, and finally all three types together. The parameters were kept consistent across experiments. Results from the experiments are presented in **Figure 4**.

**Figure 4:**
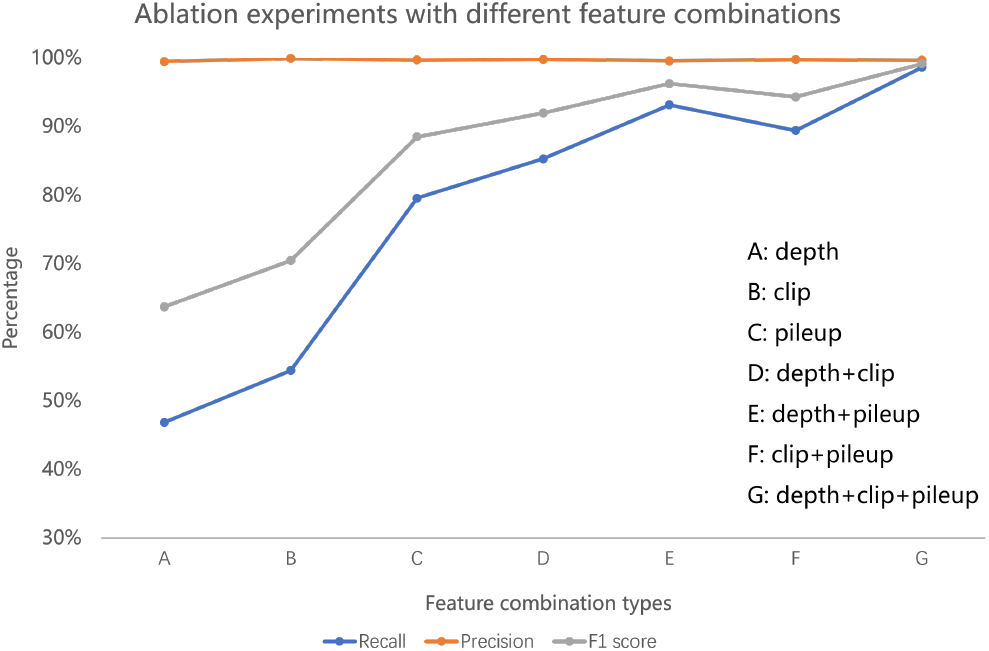
Ablation experiments with different feature on Simulated CHM13 HiFi dataset.

From the results of the ablation experiments, the precision of misassembly detection across various features approaches 100%, owing to our implementation of a rigorous filtering step for candidate assembly errors to ensure precision. However, recall exhibits notable variation. When using LRMD with a single feature, recall falls below 80%, resulting in a significant number of missed misassemblies. Integrating multiple features substantially improves recall, increasing from 40% to 60% with a single feature to over 95% with three features. Each feature contributes differently to recall, with pileup information being the most influential. In conclusion, the ablation experiment results underscore the critical importance of integrating multiple features to enhance recall in the process.

### 3.3 Benchmark on real datasets

We utilized two PacBio HiFi datasets (*O. sativa* and CHM13) and two Nanopore datasets (*O. sativa* and CHM13) to assess the performance of LRMD. The PacBio HiFi datasets were assembled using Flye and hifiasm, while the Nanopore datasets were assembled using Flye and wtdbg2 (refer to **Methods**). LRMD was applied to these eight assemblies, as detailed in **Table 3** and **Table 4**.

**Table 3:** Statistics of the assemblies.

| Data | Assembly | Contigs | Length (Gb) | Max (Mb) | N50 (Mb) |
| --- | --- | --- | --- | --- | --- |
| O. sativa, ONT | flye | 86 | 0.39 | 43.9 | 16.8 |
|  | wtdbg2 | 10437 | 0.83 | 4.2 | 0.2 |
| CHM13, ONT | flye | 680 | 2.86 | 110.0 | 51.9 |
|  | wtdbg2 | 9421 | 3.05 | 31.9 | 7.3 |
| O. sativa, HiFi | flye | 77 | 0.38 | 33.7 | 14.1 |
|  | hifiasm | 137 | 0.39 | 39.5 | 30.4 |
| CHM13, HiFi | flye | 1451 | 3.00 | 114.7 | 40.1 |
|  | hifiasm | 385 | 3.05 | 201.1 | 87.7 |

Max indicates the maximum contig length. N50 is the length of the shortest contig for which longer and equal length contigs cover at least 50 of assembly size.

**Table 4:** Misassembly and base quality evaluated by the tools on the real assemblies.

| Data | Assembly | Misassembly |  |  |  | QV |  |  |  |
| --- | --- | --- | --- | --- | --- | --- | --- | --- | --- |
|  |  | GAEP<sup>1</sup> | CRAQ | Inspector | LRMD | QUAST | Inspector | LRMD | QUAST<sup>2</sup> |
| O. sativa, ONT | flye | 337 | 1349 | 3 | 31 | 51/887 | 47.06 | 28.02 | 19.93 |
|  | wtdbg2 | 1095 | 19030 | 331 | 5109 | 603/33620 | 19.96 | 23.15 | 13.79 |
| CHM13, ONT | flye | - | 1349 | 70 | 322 | 351/845 | 43.32 | 41.93 | 29.37 |
|  | wtdbg2 | 5449 | 19030 | 1279 | 5214 | 1996/12590 | 26.43 | 33.13 | 23.81 |
| O. sativa, HiFi | flye | - | 160 | 2 | 14 | 39/12 | 64.62 | 61.72 | 43.75 |
|  | hifiasm | 19 | 282 | 17 | 48 | 105/56 | 53.06 | 56.09 | 41.48 |
| CHM13, HiFi | flye | - | 2921 | 11 | 133 | 121/95 | 56.88 | 57.78 | 47.19 |
|  | hifiasm | - | 778 | 3 | 37 | 35/88 | 60.24 | 61.27 | 49.43 |

<sup>1</sup> GAEP did not yield any results within a week on four assemblies, so the corresponding results are excluded.
<sup>2</sup> QUAST is a reference-based tools. It reports two types of misassemblies: misassemblies and local misassemblies, separated by slash.

The sizes of these assemblies closely resemble their corresponding reference genomes, except for the wtdbg2 assembly of O. sativa from Nanopore data, which is notably fragmented with an N50 of contigs at only 201 Kb and over 10,000 contigs. In contrast, other assemblies appear relatively typical. Additionally, analysis of N50 values reveals that for PacBio HiFi data, hifiasm produces more contiguous assemblies compared to Flye. Conversely, for Nanopore datasets, Flye’s assemblies demonstrate superior contiguity compared to those generated by wtdbg2.

LRMD can report the base quality (QV) of assemblies (see **Methods**). As shown in **Table 4**, for the assemblies of PacBio HiFi datasets, the QV values evaluated by LRMD are 56.09-61.72, while for the assemblies of Nanopore datases, the QV values evaluated by LRMD are 23.15-41.93. Compared with the QV results evaluated by QUAST (**Table 4** ), LRMD slightly overestimated the base quality. But QV reported by LRMD are consistent with those by QUAST, that is, the assembly with higher QV evaluated by LRMD have higher QV output by QUAST.

LRMD evaluated assembly errors in assemblies produced by wtdbg2, reporting 5019 and 5214 misassemblies, significantly higher than those of other tools, as detailed in **Table 4**. Therefore, wtdbg2’s assemblies on these real datasets exhibit inferior performance in terms of misassemblies compared to other tools. We also utilized three reference-free tools (GAEP, CRAQ, Inspector) and the reference-based tool QUAST on these datasets. All the tools report fewer misassemblies on HiFi assemblies compared to Nanopore assemblies.

We further assessed the Jaccard coefficient (see **Methods**) to measure the similarity of misassemblies identified by GAEP, CRAG, Inspector, and LRMD relative to QUAST, as detailed in **Table 5**. QUAST reports two types of misassemblies: misassemblies and local misassemblies. We combine these for evaluation in our assessment. Compared to GAEP, CRAQ, and Inspector, LRMD achieved the highest Jaccard coefficient across all assemblies. LRMD attained higher similarity, particularly notable in the assembly of CHM13 Nanopore data by Flye, where LRMD’s similarity with QUAST (0.423) surpassed that of GAEP (0.164), CRAG (0.018), and Inspector (0.030). From **Figure 5**, LRMD identifies 56.3% of misassemblies also detected by QUAST, while QUAST identifies 62.9% of LRMD’s identified misassemblies.

**Table 5:** Jaccard coefficient between misassemblies identified by QUAST and those identified by other tools on the real assemblies.

| Data | Tool | CRAQ | Inspector | LRMD | GAEP<sup>1</sup> |
| --- | --- | --- | --- | --- | --- |
| O. sativa, ONT | flye | 0.007 | 0.001 | **0.019** | 0.015 |
|  | wtdbg2 | 0.026 | 0.012 | **0.236** | 0.027 |
| CHM13, ONT | flye | 0.018 | 0.030 | **0.423** | - |
|  | wtdbg2 | 0.022 | 0.105 | **0.348** | 0.164 |
| O. sativa, HiFi | flye | 0.013 | 0.000 | **0.026** | - |
|  | hifiasm | 0.011 | 0.000 | **0.189** | 0.040 |
| CHM13, HiFi | flye | 0.010 | 0.006 | **0.136** | - |
|  | hifiasm | 0.007 | 0.000 | **0.049** | - |

<sup>1</sup> GAEP did not yield any results within a week on four assemblies, so the corresponding results are excluded.

**Figure 5:**
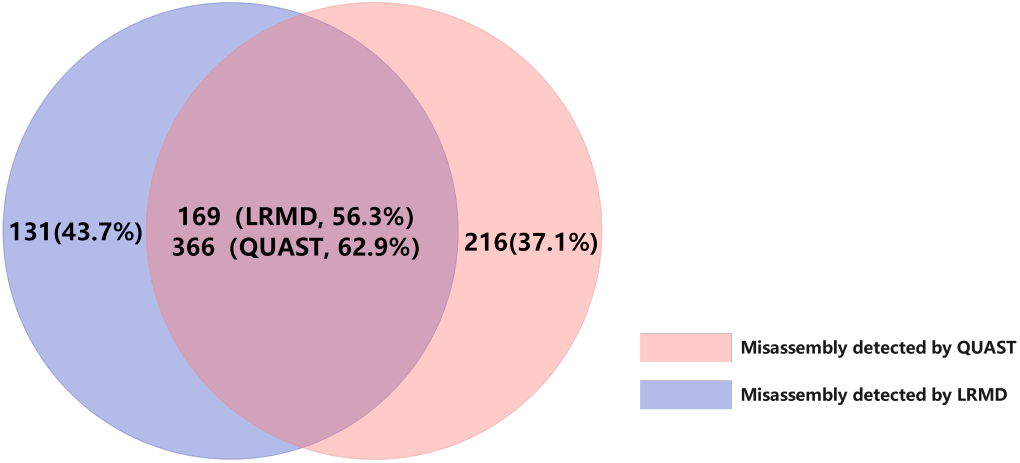
Comparison of misassembly detected by LRMD and QUAST on the assembly of CHM13 ONT data by Flye.

Additionally, the runtime and memory usage of GAEP, CRAQ, Inspector, and LRMD for evaluating each real assembly is documented in **Table 6**. They do not include the time and memory used during the alignment step. LRMD is not the fastest tool, partly because it performs multiple alignment checks to utilize various features. From **Table 6**, it can be seen that the alignment step is the performance bottleneck of the reference-free misassembly detection tools. In summary, these findings underscore LRMD’s effectiveness in evaluating real-world assemblies and detecting their associated assembly errors.

**Table 6:** CPU time and memory usage of the tools on real assemblies.

| Assembly | Tool | CPU time<br>(h) | Peak memory<br>(Gb) |
| --- | --- | --- | --- |
| O. sativa, ONT<br>flye | minimap2<sup>1</sup> | 5.6 | 61.7 |
|  | GAEP<sup>2</sup> | 1.2 | 4.2 |
|  | CRAQ | 3.0 | 107.3 |
|  | Inspector | **1.0** | **0.8** |
|  | LRMD | 1.9 | 4.8 |
| O. sativa, ONT<br>wtdbg2 | minimap2 | 8.5 | 69.7 |
|  | GAEP | **1.9** | 6.9 |
|  | CRAQ | 3.7 | 111.2 |
|  | Inspector | 2.2 | 2.0 |
|  | LRMD | 3.9 | **1.0** |
| CHM13, ONT<br>flye | minimap2 | 112.5 | 92.5 |
|  | GAEP | - | - |
|  | CRAQ | 10.8 | 178.2 |
|  | Inspector | **7.3** | **5.5** |
|  | LRMD | 13.3 | 12.3 |
| CHM13, ONT<br>wtdbg2 | minimap2 | 86.3 | 92.3 |
|  | GAEP | **8.3** | 12.2 |
|  | CRAQ | 10.3 | 168.9 |
|  | Inspector | 8.4 | 5.8 |
|  | LRMD | 15.3 | **3.9** |
| O. sativa, HiFi<br>flye | minimap2 | 4.0 | 50.9 |
|  | GAEP | - | - |
|  | CRAQ | 1.1 | 11.2 |
|  | Inspector | **0.7** | **0.8** |
|  | LRMD | 1.7 | 4.3 |
| O. sativa, HiFi<br>hifiasm | minimap2 | 4.6 | 49.4 |
|  | GAEP | **0.2** | 0.8 |
|  | CRAQ | 0.9 | 11.1 |
|  | Inspector | 0.7 | **0.9** |
|  | LRMD | 1.7 | 4.3 |
| CHM13, HiFi<br>flye | minimap2 | 18.0 | 78.9 |
|  | GAEP | - | - |
|  | CRAQ | 9.7 | 55.1 |
|  | Inspector | **4.9** | **5.8** |
|  | LRMD | 13.0 | 12.7 |
| CHM13, HiFi<br>hifiasm | minimap2 | 17.3 | 75.1 |
|  | GAEP | - | - |
|  | CRAQ | 7.5 | 54.7 |
|  | Inspector | **4.9** | **5.8** |
|  | LRMD | 13.0 | 12.7 |

<sup>1</sup> minimap2 is used for aligning long reads to assemblies.
<sup>2</sup> GAEP did not yield any results within a week on four assemblies, so the corresponding results are excluded.

## 4 DISCUSSION

High-quality assembly is crucial for downstream omics analysis. Misassembly detection is a necessary step to identify potential errors. Reference-free methods for misassembly detection, which do not rely on a reference genome, offer promising applications. In this study, we developed LRMD, a reference-free misassembly detection method that utilizes multiple features extracted from read-vs-assembly alignments. This approach significantly enhances detection recall. A candidate misassembly filtering algorithm is employed to eliminate false positives and improve method precision. Compared to existing reference-free tools, LRMD demonstrates superior recall, achieving the highest F1-Score on simulated datasets. On real datasets, LRMD shows more consistent results with the reference-based tool QUAST than other reference-free tools. By enhancing detection recall, LRMD provides valuable guidance for further improving assembly quality. Building on this foundation, we will further develop tools to correct misassemblies.

Currently, LRMD is constrained by the length of reads, making it difficult to detect misassemblies that exceed the length of long reads. This limitation is a drawback of reference-free detection tools. Hi-C sequencing reads offer long-range information about the entire chromosome structure [35–37]. Integrating Hi-C sequencing reads with reference-free tools could enhance the detection of more complex misassemblies. Effectively leveraging Hi-C sequence data is a key focus of our future research. While LRMD significantly enhances misassembly recall, it does not classify misassembly types or correct assemblies accordingly. Addressing these aspects will be a priority for our future improvements to LRMD.

## ACKNOWLEDGMENTS

This work was supported in part by the National Natural Science Foundation of China under Grants (Nos. 62350004, 62332020), the Project of Xiangjiang Laboratory (No. 23XJ01011), Fundamental Research Funds for the Central Universities of Central South University (2021zzts0208). We are grateful for resources from the High-Performance Computing Center of Central South University.

## SUPPLEMENTARY

### A SIMULATED READS GENERATION

1. Nanopore data simulation command: `pbsim -strategy wgs -method qshmm -qshmm $model -depth $coverage -genome $genome -prefix $prefix,` model selected `QSHMM-ONT.model`, depth of 40X.
2. The pbsim3 simulator cannot directly generate simulated high-precision PacBio HiFi reads. This paper first uses pbsim3 to simulate the multi-pass sequencing of sub-reads according to the method described in the pbsim3 article, and then uses the bam data generated by pbsim3 as input to generate the final simulated PacBio HiFi reads using the default parameters of We combined it with pbccs, following the method recommended by PBSIM3, to generate HiFi reads. The commands for simulating PacBio HiFi reads are as follows:
  a. `pbsim -strategy wgs -method qshmm -qshmm $model -depth $coverage -genome $fa_in -prefix $prefix -pass-num $pass_num`, where pass_num is 10 passes, model is `QSHMM-RSII.model`, and the depth is 40X;
  b. parallel -joblog samtools.log -j$threads “samtools view -bS $out_dir/{1}.sam > $out_-dir/{1}.bam && rm $out_dir/{1}.sam” ::: $sam_ls[@];
  c. parallel -joblog ccs.log -j$threads “ccs $out_dir/{1}.bam $out_dir/{1}.fastq.gz -j 2 && rm $out_dir/{1}.bam” ::: $sam_ls[@].
3. In addition, to reproduce the results in GAEP, we used pbsim2 (https://github.com/yukiteruono/pbsim2) to generate Pacbio CLR reads for the GRCH38 simulated reference sequence, with the same parameters as in the GAEP `paper: -depth 50 -length-min 5000 -length-max 50,000 -length-mean 20,000.`

## REFERENCES

[1] Yunhao Wang, Yue Zhao, Audrey Bollas, Yuru Wang, and Kin Fai Au. 2021. Nanopore Sequencing Technology, Bioinformatics and Applications. Nature Biotechnology 39, 11 (Nov. 2021), 1348–1365. 10.1038/s41587-021-01108-x

[2] Glennis A. Logsdon, Mitchell R. Vollger, and Evan E. Eichler. 2020. Long-Read Human Genome Sequencing and Its Applications. Nature Reviews Genetics 21, 10 (Oct. 2020), 597–614. 10.1038/s41576-020-0236-x

[3] Heng Li. 2016. Minimap and Miniasm: Fast Mapping and de Novo Assembly for Noisy Long Sequences. Bioinformatics 32, 14 (July 2016), 2103–2110. 10.1093/bioinformatics/btw152

[4] Sergey Koren, Brian P. Walenz, Konstantin Berlin, Jason R. Miller, Nicholas H. Bergman, and Adam M. Phillippy. 2017. Canu: Scalable and Accurate Long-Read Assembly via Adaptive k-Mer Weighting and Repeat Separation. Genome Research 27, 5 (Jan. 2017), 722–736. 10.1101/gr.215087.116

[5] Jue Ruan and Heng Li. 2020. Fast and Accurate Long-Read Assembly with Wtdbg2. Nature Methods 17, 2 (Feb. 2020), 155–158. 10.1038/s41592-019-0669-3

[6] Mikhail Kolmogorov, Jeffrey Yuan, Yu Lin, and Pavel A. Pevzner. 2019. Assembly of Long, Error-Prone Reads Using Repeat Graphs. Nature Biotechnology 37, 5 (May 2019), 540–546. 10.1038/s41587-019-0072-8xs

[7] Haoyu Cheng, Gregory T. Concepcion, Xiaowen Feng, Haowen Zhang, and Heng Li. 2021. Haplotype-Resolved de Novo Assembly Using Phased Assembly Graphs with Hifiasm. Nature Methods 18, 2 (Feb. 2021), 170–175. 10.1038/s41592-020-01056-5

[8] Mikko Rautiainen, Sergey Nurk, Brian P. Walenz, Glennis A. Logsdon, David Porubsky, Arang Rhie, Evan E. Eichler, Adam M. Phillippy, and Sergey Koren. 2023. Telomere-to-Telomere Assembly of Diploid Chromosomes with Verkko. Nature Biotechnology 41, 10 (Oct. 2023), 1474–1482. 10.1038/s41587-023-01662-6

[9] Heng Li and Richard Durbin. 2024. Genome Assembly in the Telomere-to-Telomere Era. Nature Reviews Genetics (April 2024), 1–13. 10.1038/s41576-024-00718-w

[10] Yanqing Sun, Lianguang Shang, Qian-Hao Zhu, Longjiang Fan, and Longbiao Guo. 2022. Twenty Years of Plant Genome Sequencing: Achievements and Challenges. Trends in Plant Science 27, 4 (April 2022), 391–401. 10.1016/j.tplants.2021.10.006

[11] Arang Rhie, Shane A. McCarthy, Olivier Fedrigo, Joana Damas, Giulio Formenti, Sergey Koren, Marcela Uliano-Silva, William Chow, Arkarachai Fungtammasan, Juwan Kim, Chul Lee, Byung June Ko, Mark Chaisson, Gregory L. Gedman, Lindsey J. Cantin, Francoise Thibaud-Nissen, Leanne Haggerty, Iliana Bista, Michelle Smith, Bettina Haase, Jacquelyn Mountcastle, Sylke Winkler, Sadye Paez, Jason Howard, Sonja C. Vernes, Tanya M. Lama, Frank Grutzner, Wesley C. Warren, Christopher N. Balakrishnan, Dave Burt, Julia M. George, Matthew T. Biegler, David Iorns, Andrew Digby, Daryl Eason, Bruce Robertson, Taylor Edwards, Mark Wilkinson, George Turner, Axel Meyer, Andreas F. Kautt, Paolo Franchini, H. William Detrich, Hannes Svardal, Maximilian Wagner, Gavin J. P. Naylor, Martin Pippel, Milan Malinsky, Mark Mooney, Maria Simbirsky, Brett T. Hannigan, Trevor Pesout, Marlys Houck, Ann Misuraca, Sarah B. Kingan, Richard Hall, Zev Kronenberg, Ivan Sović, Christopher Dunn, Zemin Ning, Alex Hastie, Joyce Lee, Siddarth Selvaraj, Richard E. Green, Nicholas H. Putnam, Ivo Gut, Jay Ghurye, Erik Garrison, Ying Sims, Joanna Collins, Sarah Pelan, James Torrance, Alan Tracey, Jonathan Wood, Robel E. Dagnew, Dengfeng Guan, Sarah E. London, David F. Clayton, Claudio V. Mello, Samantha R. Friedrich, Peter V. Lovell, Ekaterina Osipova, Farooq O. Al-Ajli, Simona Secomandi, Heebal Kim, Constantina Theofanopoulou, Michael Hiller, Yang Zhou, Robert S. Harris, Kateryna D. Makova, Paul Medvedev, Jinna Hoffman, Patrick Masterson, Karen Clark, Fergal Martin, Kevin Howe, Paul Flicek, Brian P. Walenz, Woori Kwak, Hiram Clawson, Mark Diekhans, Luis Nassar, Benedict Paten, Robert H. S. Kraus, Andrew J. Crawford, M. Thomas P. Gilbert, Guojie Zhang, Byrappa Venkatesh, Robert W. Murphy, Klaus-Peter Koepfli, Beth Shapiro, Warren E. Johnson, Federica Di Palma, Tomas Marques-Bonet, Emma C. Teeling, Tandy Warnow, Jennifer Marshall Graves, Oliver A. Ryder, David Haussler, Stephen J. O’Brien, Jonas Korlach, Harris A. Lewin, Kerstin Howe, Eugene W. Myers, Richard Durbin, Adam M. Phillippy, and Erich D. Jarvis. 2021. Towards Complete and Error-Free Genome Assemblies of All Vertebrate Species. Nature 592, 7856 (April 2021), 737–746. 10.1038/s41586-021-03451-0

[12] Ole K Tørresen, Bastiaan Star, Pablo Mier, Miguel A Andrade-Navarro, Alex Bateman, Patryk Jarnot, Aleksandra Gruca, Marcin Grynberg, Andrey V Kajava, Vasilis J Promponas, Maria Anisimova, Kjetill S Jakobsen, and Dirk Linke. 2019. Tandem Repeats Lead to Sequence Assembly Errors and Impose Multi-Level Challenges for Genome and Protein Databases. Nucleic Acids Research 47, 21 (Dec. 2019), 10994–11006. 10.1093/nar/gkz841

[13] Mick Watson and Amanda Warr. 2019. Errors in Long-Read Assemblies Can Critically Affect Protein Prediction. Nature Biotechnology 37, 2 (Feb. 2019), 124– 126. 10.1038/s41587-018-0004-z

[14] James F. Denton, Jose Lugo-Martinez, Abraham E. Tucker, Daniel R. Schrider, Wesley C. Warren, and Matthew W. Hahn. 2014 –12–4–. Extensive Error in the Number of Genes Inferred from Draft Genome Assemblies. PLOS Computational Biology 10, 12 (2014–12–4–), e1003998. 10.1371/journal.pcbi.1003998

[15] Adam Thrash, Federico Hoffmann, and Andy Perkins. 2020. Toward a More Holistic Method of Genome Assembly Assessment. BMC Bioinformatics 21, 4 (July 2020), 249. 10.1186/s12859-020-3382-4

[16] Elena Espinosa, Rocio Bautista, Rafael Larrosa, and Oscar Plata. 2024. Advancements in Long-Read Genome Sequencing Technologies and Algorithms. Genomics 116, 3 (May 2024), 110842. 10.1016/j.ygeno.2024.110842

[17] Quanyu Chen, Chentao Yang, Guojie Zhang, and Dongya Wu. 2024. GCI: a continuity inspector for complete genome assembly. bioRxiv (2024). 10.1101/2024.04.06.588431 arXiv:https://www.biorxiv.org/content/early/2024/04/09/2024.04.06.588431.full.pdf

[18] Mosè Manni Matthew R Berkeley, Mathieu Seppey, Felipe A Simão, and Evgeny M Zdobnov. 2021. BUSCO Update: Novel and Streamlined Workflows along with Broader and Deeper Phylogenetic Coverage for Scoring of Eukaryotic, Prokaryotic, and Viral Genomes. Molecular Biology and Evolution 38, 10 (Oct. 2021), 4647–4654. 10.1093/molbev/msab199

[19] Wen-Biao Jiao, Gonzalo Garcia Accinelli, Benjamin Hartwig, Christiane Kiefer, David Baker, Edouard Severing, Eva-Maria Willing, Mathieu Piednoel, Stefan Woetzel, Eva Madrid-Herrero, Bruno Huettel, Ulrike Hümann, Richard Reinhard, Marcus A. Koch, Daniel Swan, Bernardo Clavijo, George Coupland, and Korbinian Schneeberger. 2017. Improving and Correcting the Contiguity of Long-Read Genome Assemblies of Three Plant Species Using Optical Mapping and Chromosome Conformation Capture Data. Genome Research 27, 5 (Jan. 2017), 778–786. 10.1101/gr.213652.116

[20] Medhat Mahmoud, Nastassia Gobet, Diana Ivette Cruz-Dávalos, Ninon Mounier, Christophe Dessimoz, and Fritz J. Sedlazeck. 2019. Structural Variant Calling: The Long and the Short of It. Genome Biology 20, 1 (Nov. 2019), 246. 10.1186/s13059-019-1828-7

[21] Tao Jiang, Yongzhuang Liu, Yue Jiang, Junyi Li, Yan Gao, Zhe Cui, Yadong Liu, Bo Liu, and Yadong Wang. 2020. Long-read-based human genomic structural variation detection with cuteSV. Genome Biology 21, 1 (Aug. 2020), 189. 10.1186/s13059-020-02107-y

[22] Moritz Smolka, Luis F. Paulin, Christopher M. Grochowski, Dominic W. Horner, Medhat Mahmoud, Sairam Behera, Ester Kalef-Ezra, Mira Gandhi, Karl Hong, Davut Pehlivan, Sonja W. Scholz, Claudia M. B. Carvalho, Christos Proukakis, and Fritz J. Sedlazeck. 2024. Detection of mosaic and population-level structural variants with Sniffles2. Nature Biotechnology (Jan. 2024). 10.1038/s41587-023-02024-y

[23] Alexey Gurevich, Vladislav Saveliev, Nikolay Vyahhi, and Glenn Tesler. 2013. QUAST: Quality Assessment Tool for Genome Assemblies. Bioinformatics 29, 8 (April 2013), 1072–1075. 10.1093/bioinformatics/btt086

[24] Alla Mikheenko, Andrey Prjibelski, Vladislav Saveliev, Dmitry Antipov, and Alexey Gurevich. 2018. Versatile Genome Assembly Evaluation with QUASTLG. Bioinformatics 34, 13 (July 2018), i142–i150. 10.1093/bioinformatics/bty266

[25] Steven L. Salzberg, Adam M. Phillippy, Aleksey Zimin, Daniela Puiu, Tanja Magoc, Sergey Koren, Todd J. Treangen, Michael C. Schatz, Arthur L. Delcher, Michael Roberts, Guillaume Marçais, Mihai Pop, and James A. Yorke. 2012. GAGE: A Critical Evaluation of Genome Assemblies and Assembly Algorithms. Genome Research 22, 3 (Jan. 2012), 557–567. 10.1101/gr.131383.111

[26] Martin Hunt, Taisei Kikuchi, Mandy Sanders, Chris Newbold, Matthew Berriman, and Thomas D. Otto. 2013. REAPR: A Universal Tool for Genome Assembly Evaluation. Genome Biology 14, 5 (May 2013), R47. 10.1186/gb2013-14-5-r47

[27] Bruce J. Walker, Thomas Abeel, Terrance Shea, Margaret Priest, Amr Abouelliel, Sharadha Sakthikumar, Christina A. Cuomo, Qiandong Zeng, Jennifer Wortman, Sarah K. Young, and Ashlee M. Earl. 2014 –11–19–. Pilon: An Integrated Tool for Comprehensive Microbial Variant Detection and Genome Assembly Improvement. PLOS ONE 9, 11 (2014–11–19–), e112963. 10.1371/journal.pone.0112963

[28] Min Li, Binbin Wu, Xiaodong Yan, Junwei Luo, Yi Pan, Fang-Xiang Wu, and Jianxin Wang. 2017. PECC: Correcting Contigs Based on Paired-End Read Distribution. Computational Biology and Chemistry 69 (Aug. 2017), 178–184. 10.1016/j.compbiolchem.2017.03.012

[29] Binbin Wu, Min Li, Xingyu Liao, Junwei Luo, Fang-Xiang Wu, Yi Pan, and Jianxin Wang. 2020. MEC: Misassembly Error Correction in Contigs Based on Distribution of Paired-End Reads and Statistics of GC-Contents. IEEE/ACM Transactions on Computational Biology and Bioinformatics 17, 3 (May 2020), 847– 857. 10.1109/TCBB.2018.2876855

[30] Yu Chen, Yixin Zhang, Amy Y. Wang, Min Gao, and Zechen Chong. 2021. Accurate Long-Read de Novo Assembly Evaluation with Inspector. Genome Biology 22, 1 (Nov. 2021), 312. 10.1186/s13059-021-02527-4

[31] Kunpeng Li, Peng Xu, Jinpeng Wang, Xin Yi, and Yuannian Jiao. 2023. Identification of Errors in Draft Genome Assemblies at Single-Nucleotide Resolution for Quality Assessment and Improvement. Nature Communications 14, 1 (Oct. 2023), 6556. 10.1038/s41467-023-42336-w

[32] Yong Zhang, Hong-Wei Lu, and Jue Ruan. 2023. GAEP: A Comprehensive Genome Assembly Evaluating Pipeline. Journal of Genetics and Genomics 50, 10 (Oct. 2023), 747–754. 10.1016/j.jgg.2023.05.009

[33] Heng Li. 2018. Minimap2: Pairwise Alignment for Nucleotide Sequences. Bioinformatics 34, 18 (Sept. 2018), 3094–3100. 10.1093/bioinformatics/bty191

[34] Yukiteru Ono, Michiaki Hamada, and Kiyoshi Asai. 2022. PBSIM3: A Simulator for All Types of PacBio and ONT Long Reads. NAR Genomics and Bioinformatics 4, 4 (Dec. 2022), qac092. 10.1093/nargab/lqac092

[35] Da Lin, Ping Hong, Siheng Zhang, Weize Xu, Muhammad Jamal, Keji Yan, Yingying Lei, Liang Li, Yijun Ruan, Zhen F. Fu, Guoliang Li, and Gang Cao. 2018. Digestion-ligation-only Hi-C is an efficient and cost-effective method for chromosome conformation capture. Nature Genetics 50, 5 (May 2018), 754–763. 10.1038/s41588-018-0111-2

[36] Hui-Su Kim, Sungwon Jeon, Changjae Kim, Yeon Kyung Kim, Yun Sung Cho, Jungeun Kim, Asta Blazyte, Andrea Manica, Semin Lee, and Jong Bhak. 2019. Chromosome-scale assembly comparison of the Korean Reference Genome KOREF from PromethION and PacBio with Hi-C mapping information. GigaScience 8, 12 (Dec. 2019), giz125. 10.1093/gigascience/giz125

[37] Zev N. Kronenberg, Arang Rhie, Sergey Koren, Gregory T. Concepcion, Paul Peluso, Katherine M. Munson, David Porubsky, Kristen Kuhn, Kathryn A. Mueller, Wai Yee Low, Stefan Hiendleder, Olivier Fedrigo, Ivan Liachko, Richard J. Hall, Adam M. Phillippy, Evan E. Eichler, John L. Williams, Timothy P. L. Smith, Erich D. Jarvis, Shawn T. Sullivan, and Sarah B. Kingan. 2021. Extended haplotype-phasing of long-read de novo genome assemblies using Hi-C. Nature Communications 12, 1 (April 2021), 1935. 10.1038/s41467-020-20536-y

